# Multidimensional diffusion MRI reveals heterogeneous microstructural remodeling associated with amyloid pathology

**DOI:** 10.64898/2026.08.26.747377

**Authors:** Pak Shing Kenneth Or, Maxime Yon, Omar Narvaez, Valeriia Sitnikova, Tarja Malm, Mustapha Bouhrara, Alejandra Sierra, Daniel Topgaard, Dan Benjamini

**Author notes:** Corresponding author: Dan Benjamini, PhD, Laboratory of Behavioral Neuroscience, National Institute on Aging, National Institutes of Health, Baltimore 21224, MD, USA.

## Abstract

Alzheimer’s disease (AD) pathology involves amyloid deposition, reactive gliosis, and localized tissue alterations that coexist within the same brain regions, creating heterogeneous microstructural environments within individual imaging voxels. Conventional diffusion MRI averages these environments into aggregate measures, potentially obscuring their distinct contributions. Frequency-dependent multidimensional MRI (ωMD-MRI) resolves distributions of water components with different diffusion length scales, anisotropies, and relaxation properties, providing sensitivity to microstructural restriction, heterogeneity, and shape-size correlations within a voxel. Whether these measurements reveal microstructural complexity associated with AD pathology remains unclear. Here, we performed ωMD-MRI on ex vivo brain specimens from approximately 8-month-old 5xFAD and wild-type mice and interpreted the imaging findings alongside complementary histology. ωMD-MRI revealed widespread but spatially nonuniform differences between 5xFAD and wild-type brains. Measurements sensitive to microstructural restriction, heterogeneity, and shape-size correlations consistently indicated greater microstructural heterogeneity in 5xFAD brains, with the most prominent differences in the hippocampal formation and major cerebral white matter tracts. Complementary qualitative histology demonstrated extensive amyloid deposition and glial activation in affected regions, while overall cytoarchitecture and myelin organization remained largely preserved. Thus, the ωMD-MRI abnormalities occurred in tissue characterized by multiple coexisting pathological and relatively preserved microstructural environments rather than widespread structural degeneration. These findings demonstrate that ωMD-MRI can reveal the spatial and microstructural heterogeneity associated with amyloid pathology and provide a more comprehensive characterization of AD-related tissue alterations.

## 1. Introduction

Alzheimer’s disease (AD) is a biologically and clinically heterogeneous disorder involving interacting processes that include protein aggregation, neuroimmune and vascular dysfunction, and synaptic, axonal, oligodendroglial, and myelin alterations^1–3^. These processes vary across individuals and brain regions and may coexist within the same tissue, potentially contributing to the imperfect relationship between molecular pathology, clinical presentation, and rates of cognitive decline. Biomarkers capable of characterizing the regional tissue consequences of this biological complexity—including microstructural remodeling not captured by gross structural measures—could complement existing biomarkers and support longitudinal evaluation of disease progression and therapeutic response.

Molecular biomarkers measured using positron emission tomography (PET), cerebrospinal fluid, and, increasingly, blood have enabled the in vivo detection of specific pathological processes associated with AD, particularly amyloid-β and tau accumulation^4–6^. However, molecular biomarker burden does not uniquely determine clinical phenotype, rate of cognitive decline, or whether and when dementia will develop, reflecting the influence of additional interacting tissue processes^7–9^. These approaches also provide different and incomplete perspectives on disease biology: PET offers regional molecular information but is costly, less accessible, and involves ionizing radiation; cerebrospinal fluid sampling is invasive; and blood biomarkers, although more accessible, do not provide spatial information about brain tissue remodeling. Complementary biomarkers are therefore needed that can be acquired non-invasively and repeatedly while characterizing the regional microstructural consequences of the broader pathological process.

Magnetic resonance imaging (MRI) provides a promising platform for complementary biomarker development because it is non-invasive, free of ionizing radiation, repeatable, and capable of mapping regional tissue changes throughout the brain. MRI is routinely used in the evaluation of cognitive impairment to assess brain structure, patterns of atrophy, vascular injury, and alternative causes of dementia^4^, and it is widely applied in experimental studies of AD pathology^10^. Quantitative methods such as diffusion tensor imaging (DTI)^11^ and relaxometry^12^ extend conventional anatomical imaging by providing indirect contrast related to tissue microstructure and composition. However, these approaches generally sample a limited contrast space and summarize complex signals using a small number of aggregate parameters. Consequently, distinct pathological processes may produce similar metric changes, while contributions from coexisting microstructural environments may remain unresolved because of relaxation differences^13^, diffusion-time or diffusion-frequency dependence, and the confounding of microscopic anisotropy with orientation dispersion^14^. The variable sensitivity of conventional diffusion measurements in heavily affected regions such as the cortex^15^ and hippocampus^16,17^ illustrates the challenge of representing heterogeneous AD-associated tissue remodeling using low-dimensional MRI metrics.

Multidimensional MRI (MD-MRI) offers a strategy for addressing these limitations by jointly encoding multiple MR contrast dimensions and quantifying their joint distributions rather than examining each dimension independently^18,19^. This approach can distinguish otherwise overlapping signal components by their combined diffusion and relaxation properties, providing complementary descriptions of coexisting microstructural environments without assigning them to predefined biological substrates. Although MD-MRI is a comparatively recent framework, it has been successfully applied across diverse biological settings, ranging from characterization of healthy neural microstructure^20^ to studies of inflammation^21^, placental dysfunction^22^, tumors^23^, and AD-related pathology^24^.

The specific MD-MRI approach used here is frequency-dependent diffusion-relaxation distribution MRI (ωMD-MRI)^25^, which combines tensor-valued encoding^26^, diffusion frequency encoding^27^ and quantitative relaxometry^28^. By jointly encoding diffusion-frequency dependence, microscopic diffusion anisotropy, and relaxation properties, ωMD-MRI helps distinguish signal components that overlap in conventional measurements^29–32^ and has been shown to improve the reproducibility of derived microstructural metrics^33^. Its data-driven inversion framework estimates multidimensional signal distributions without imposing predefined tissue compartments^34,35^. By characterizing components with differing diffusion and relaxation properties, ωMD-MRI may reveal heterogeneous tissue remodeling that is averaged within conventional diffusion measurements^36^.

In the present study, we used the 5xFAD mouse as an experimental system to determine whether ωMD-MRI can resolve heterogeneous tissue remodeling associated with extensive amyloid pathology. The model coexpresses five familial AD mutations^37^ and develops widespread amyloid-β deposition, particularly within the cortex and hippocampal formation^37–39^, together with pronounced neuroinflammatory and other tissue alterations. We evaluated ωMD-MRI at high spatial resolution in ex vivo brain specimens from approximately 8-month-old 5xFAD and wild-type mice and interpreted the multidimensional imaging measurements alongside complementary histological analyses. We hypothesized that ωMD-MRI would reveal spatially and microstructurally heterogeneous alterations across gray and white matter by resolving complementary signal components that may be obscured when diffusion behavior is summarized using conventional aggregate metrics.

## 2. Methods

### 2.1. Sample preparation

Eight female mice from the 5xFAD strain, four of which are of the wild type (WT) and the other four of the AD transgenic type, were perfused at approximately 8 months old (Supp Table 1) with 0.9% saline for 2 min at 5 mL/min and then phosphate-buffered 4% paraformaldehyde solution for 20 min at 5 mL/min. Brains were extracted and fit in 10 mm NMR tubes, stored in phosphate-buffered 2% paraformaldehyde solution at 4 °C until imaging. Ethics approval was obtained from the Animal Ethics Committee of the Province Government of Southern Finland conforming to the guidelines set by the European Community Council Directives 2010/63/EEC.

**Table 1.** Regions of interest (ROIs) delineated in data with atlas.

| ROI type | Regions of interest |
| --- | --- |
| Gray matter | Cerebellar cortex, Cortical plate, Cortical subplate, Hypothalamus, Medulla, Pallidum, Pons, Striatum, Thalamus, Midbrain (Behavioral state related), Midbrain (Motor related), Midbrain (Sensory related), Hippocampus proper, Dentate gyrus, Entorhinal cortex, Subicular complex |
| White matter | Cerebellum related fiber tracts, Lateral forebrain bundle system, Medial forebrain bundle system, Supra callosal cerebral white matter |

### 2.2. MRI acquisition and processing

All acquisitions were performed on a Bruker Avance-Neo 11.7 T spectrometer (equipped with a MIC-5 probe, 3 T/m maximum gradient amplitude) with the preclinical imaging software ParaVision 360. For the *ω*MD-MRI acquisitions, we used an EPI readout and reversed phase encoding with a partial Fourier factor of 6/8 at 150 µm isotropic resolution, resulting in a total experiment time of ∼24 hours per sample. The *ω*MD-MRI acquisition consists of a series of 352 MRI acquisitions simultaneously varying echo time (*τ*_E_), recovery time (*τ*_R_) and the tensor-valued encoding spectrum **b**(*ω*). Figure 1 plots various quantities calculated from **b**(*ω*), such as its trace *b* and normalized anisotropy *b*_Δ_, and, *τ*_E_ and *τ*_R_ of the acquisition scheme. To robustly sample the acquisition space, the protocol was designed to measure at a wide range of acquisition parameters, resulting in a *b* range between 3·10^7^ and 8·10^9^ sm^-2^, *τ*_E_ between 9.4 and 49.4 ms, and *τ*_R_ between 0.8 and 3.5 s. Frequency encoding range was controlled by changing the modulation order of the double rotation of the *q* vector encoding waveform^40^, which was chosen to be between orders 0 and 2^41^. In addition to the *ω*MD-MRI acquisition, standard 3D FLASH scans at 50 µm isotropic resolution were acquired for each sample. All acquisitions were performed at 298 K.

**Figure 1.**
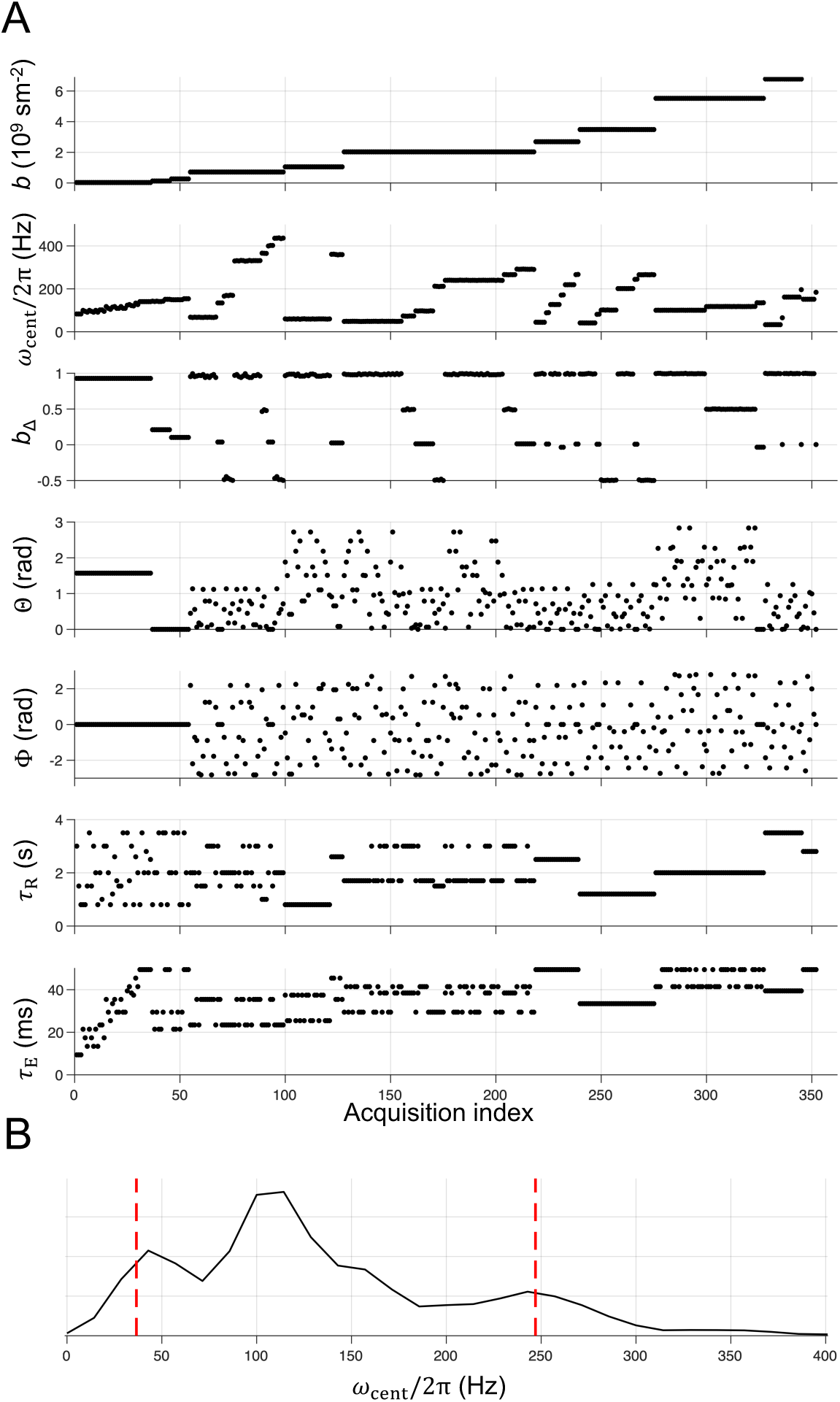
Details of the MRI acquisition scheme. (A) Acquisition parameters of the multidimensional MRI protocol used in this study, including the trace *b*, centroid frequency *ω*_cent_/2π, normalised anisotropy *b*_Δ_ and orientation (Θ, Φ) calculated from the tensor-valued diffusion encoding spectrum **b**(*ω*), recovery times *τ*_R_ and echo time *τ*_E_. (B) Histogram of *ω*_cent_/2π, where red lines depict the 10^th^ and 90^th^ percentiles at which the frequency dependence is evaluated.

The acquired *ω*MD-MRI images were preprocessed with MP-PCA denoising^42^, Gibbs ringing removal on MRtrix3^43^ and EPI distortion correction using TopUp from FSL^44^. After preprocessing, we use a Monte bootstrap inversion^35,45^ procedure to fit the acquired signal *S*(**b**(*ω*), *τ*_R_, *τ*_E_) within each voxel to the following multi-component equation^25^

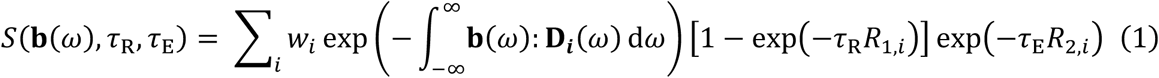

where *w_i_* is the weight of each component *i*, **D*_i_***(*ω*) the tensor-valued diffusion spectra^46,47^ in the laboratory frame of reference (assumed axially-symmetric), and *R*_1,*i*_ and *R*_2,*i*_ the longitudinal and transverse relaxation rates of each component respectively. *ω* is the diffusion encoding frequency. It is worth clarifying that although Fig. 1 shows quantities calculated from **b**(*ω*) such as its trace *b*, normalized anisotropy *b*_Δ_ and centroid frequency^40,48^ *ω*_cent_/2π, the fitting is performed using **b**(*ω*) following Eq. 1 and the calculated quantities are only displayed for bookkeeping.

In short, a multi-component, multidimensional solution in the **D**(*ω*)-*R*_1_-*R*_2_ space is obtained in each bootstrap. In this work, the maximum number of output components was limited to 10, and number of bootstrap repetitions was set to 100.

In the framework **D*_i_***(*ω*) is assumed to be axially-symmetric, such that in the principal axis system (PAS) of the tensor two of the three eigenvalues are equivalent and parameterized by Lorentzians with the zero-frequency axial diffusivity *D*_A,i_ and radial diffusivity *D*_R,i_^25,49^

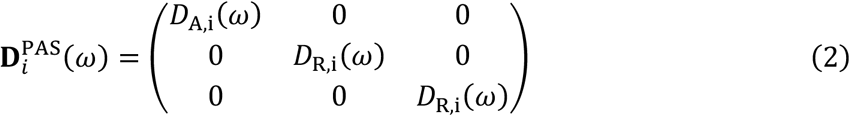

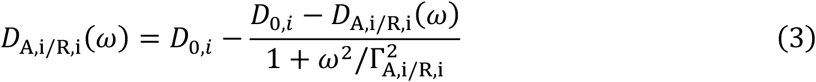

where Γ_A,i_ and Γ_R,i_ are the axial and radial transition frequencies, and *D*_0,*i*_ is the high-frequency isotropic diffusivity. The isotropic diffusivity *D*_iso_(*ω*) and normalized diffusion anisotropy *D*_Δ_(*ω*) can then be obtained^48^

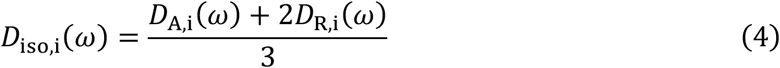

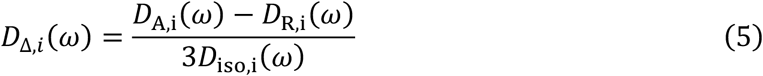

Collecting solutions over the bootstraps gives a distribution of solutions for each voxel. To facilitate visualization of these results across the entire sample, the distribution in each voxel can be characterized by statistical descriptors^18^ such as the mean E[*x*] and variance V[*x*] along any singular dimension of the solution space, or covariance C[*x*, *y*] along any two dimensions at a given *ω*. The same can also be done for an arbitrarily defined sub-space of the solution space, or ‘bin’. The following boundaries are used to define the bins^18^

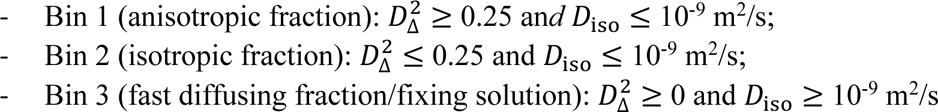

The fraction of components belonging to the *N*^th^ bin within a bootstrap *f*_bin_ *_N_* is given by

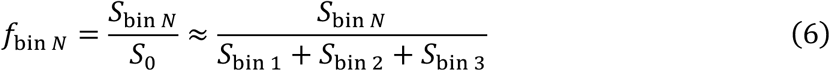

where *S*_bin_ *_N_* is the sum of the weights of the components belonging to the *N*^th^ bin, and *S*_0_ is the sum of the weights of all the components.

By evaluating 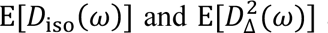 at two *ω* values (*ω*_min_ and *ω*_max_), the linear rate of change of the metrics^50,51^ Δ*_ω_*_/2*π*_E[*x*] can be obtained by

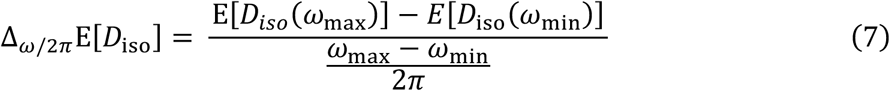

and similarly, for V[*x*] and C[*x*, *y*] metrics, and for 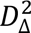. These metrics will be referred to as Δ*_ω_*_/2*π*_ metrics^49^.

ROIs were defined by registration to the Turone Mouse Brain Template and Atlas (NITRC – www.nitrc.org) with Advanced Normalization Tools^52^ (ANTs version 2.5.1, http://stnava.github.io/ANTs/, accessed on 22 April 2024) (Fig 2). The choice of ROIs can be found in Table 1.

**Figure 2.**
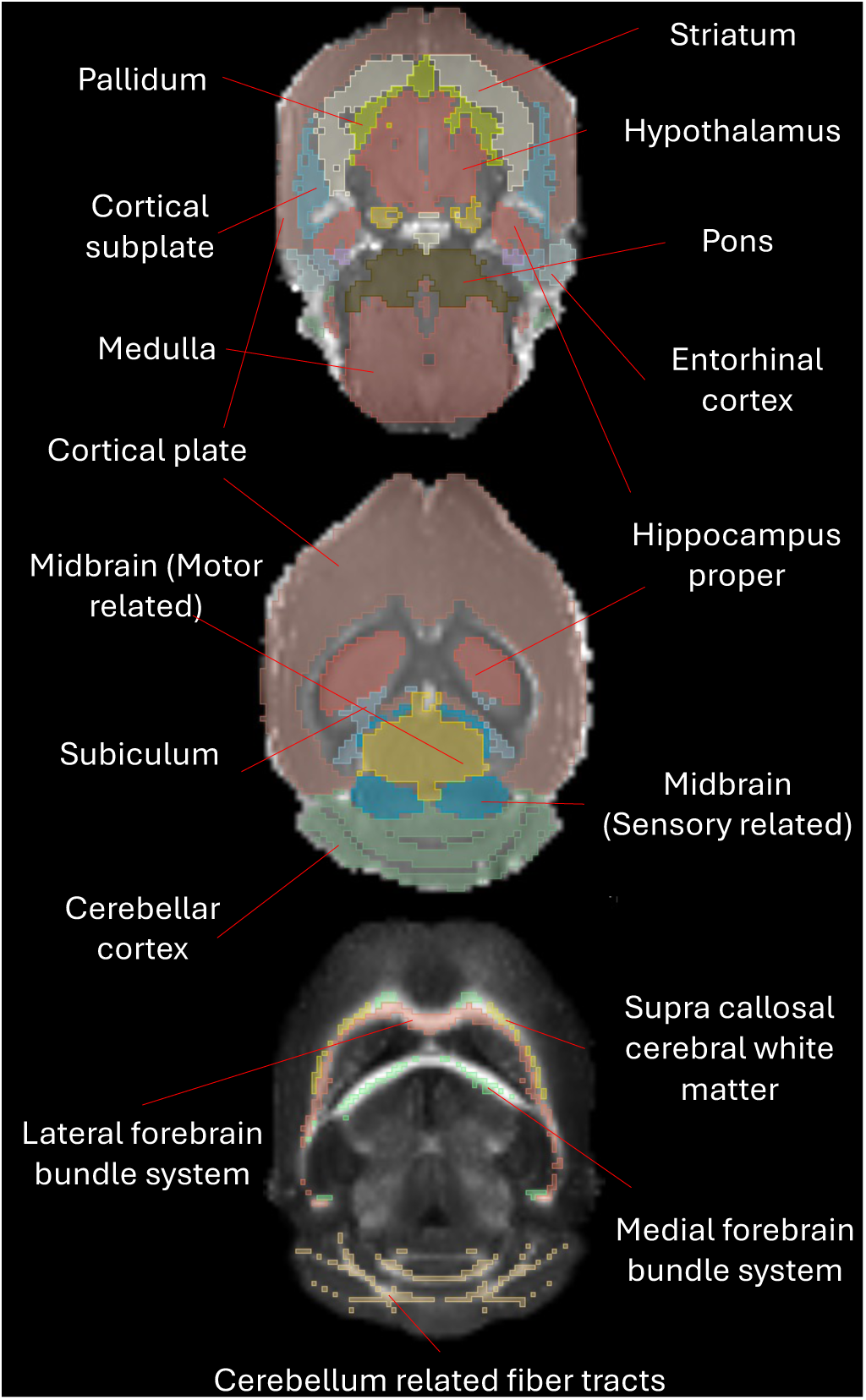
Select regions of interests (ROI) overlayed on an *S*_0_ signal intensity image and 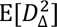 anisotropy parameter map.

### 2.3. Visualization and statistical analysis of group differences

Qualitative and quantitative methods were used to investigate the differences between the two groups of *ex vivo* mouse brain samples.

#### 2.3.1. ROI-averaged solution distribution

As described above, the Monte Carlo bootstrap inversion method^35^ yields a multi-component solution in the **D**(*ω*) − *R*_1_ − *R*_2_solution space for each bootstrap in each voxel, and collecting solutions over the bootstrap replicates gives a distribution of solutions within each voxel. Solution components can further be collected over voxels within an ROI to give an ROI-averaged solution distribution. Limited by the lower resolution of the *ω*MD-MRI scans relative to the atlas (150 µm vs 60 µm isotropic), coarser ROIs were chosen. The solutions collected from different voxels were normalized by *S*_0_ (Eq 6) to remove the effect of solution weight variation from extrinsic factors such as coil spatial sensitivity. The ROI-averaged distributions had a large number of solution components (∼1000 × number of voxels in the ROI), removing the need for Gaussian smoothing and was more feasible to inspect visually, but introduces spatial ambiguity in the interpretation of the resulting distribution. The multidimensional solution distributions were projected onto multiple two-dimensional (2D) axes for visual inspection (Fig 3).

**Figure 3.**
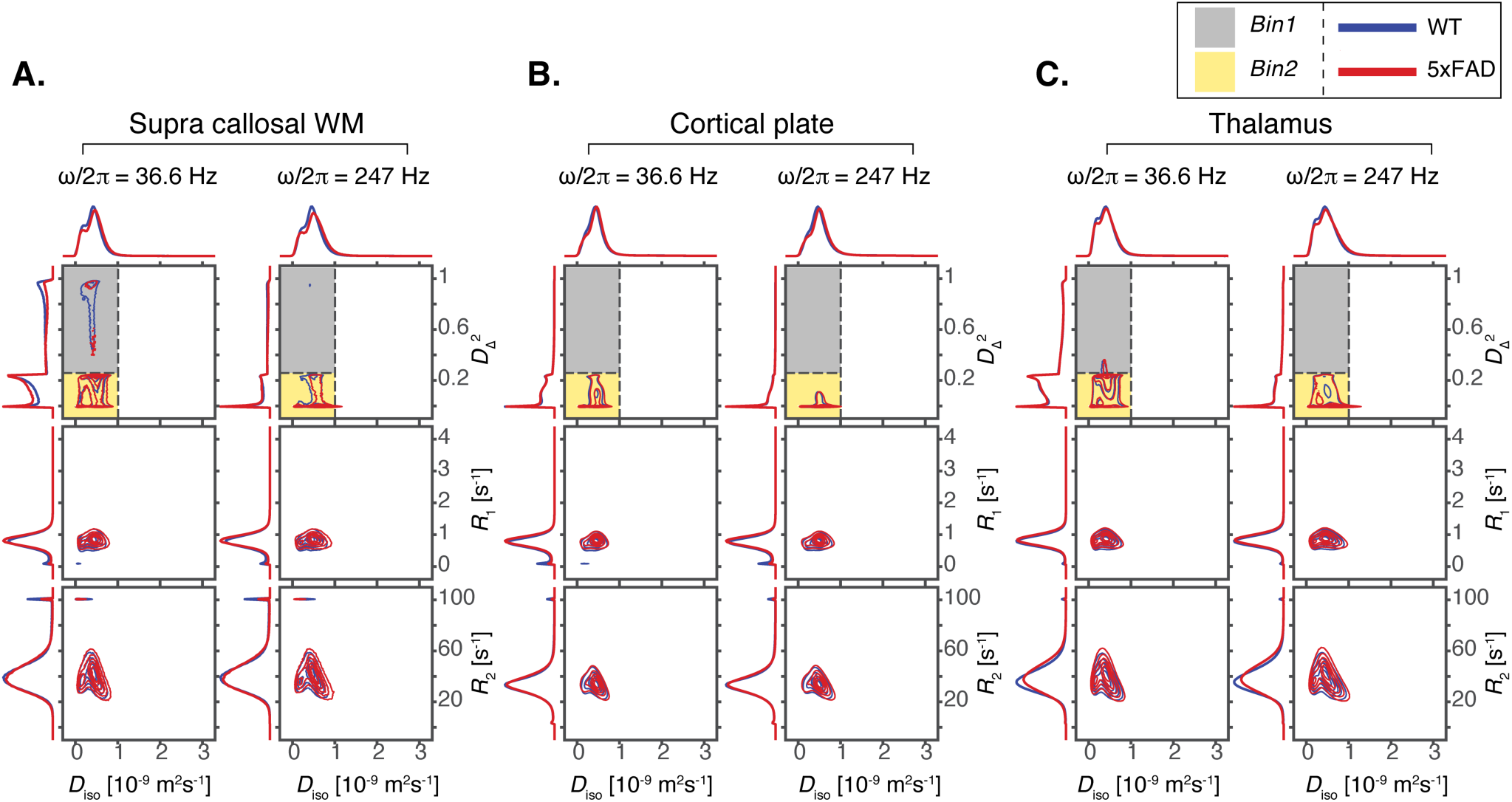
2D projections along various combinations of dimensions, including isotropic diffusivity *D*_iso_, squared normalized anisotropy 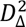, longitudinal relaxation rate *R*_1_ and transverse relaxation rate *R*_2_, of ROI-specific solution distributions (averaged over samples in the same group) in (A) supra callosal WM, (B) cortical plate and (C) thalamus. Within each subplot, the left column shows the projections at a low diffusion frequency (*ω*/2*π* = 36.6 Hz) while the right column shows that at a high diffusion frequency (*ω*/2*π* = 247 Hz). Each column shows 3 panels, which from top to bottom are the ROI-averaged solution projected to the 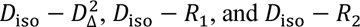 dimensions. On the top panels, bin region definitions are shaded in gray for bin 1 (high anisotropy, low diffusivity) and in yellow for bin 2 (low anisotropy and diffusivity). At the exterior of the panels show 1D projections of the distributions. Shown in blue are the ROI-specific solution distribution projections averaged over WT samples, while in red are that for AD samples.

#### 2.3.2. Group-averaged metric maps

To facilitate the examination of differences in the metric maps, the data from each of the eight subjects were registered to a common space, created with the ANTs^52^ functionality ‘antsMultivariateTemplateConstruction’. Within this space, each parameter map was averaged separately for each group (WT and AD), and voxel-wise percent change maps from WT to AD, calculated by

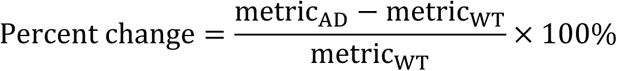

were computed to identify systematic differences.

#### 2.3.3 Linear regression of ROI-average parameters

The voxel-wise values of each metric were averaged within each ROI to generate an ROI-level metric value. For bin-resolved metrics, the ROI-average was weighted by the corresponding bin fraction to ensure voxels with a significant bin fraction of the specific bin contributes more to the estimation of the bin-resolved metric. Using these ROI-averaged metrics, we investigate group differences of each ROI-averaged metric by linear regression of the Z-score standardized ROI-averaged metrics (standardized separately in each metric and ROI), giving a *p*-value and beta coefficient (as effect size) for each. False discovery rate correction was not applied due to the small sample size.

### 2.4. Histology

After *ex vivo* imaging, the brains were washed twice for 25 min in 0.1 M phosphate buffer pH 7.4 and then cryoprotected for 48 h in a solution containing 20% glycerol in 0.02 M potassium phosphate-buffered saline pH 7.4. After cryoprotection, the brains were frozen in dry ice and stored at −70 °C until further processing. The brains were horizontally sectioned into 20-μm thick section and 1:5 series using sliding microtome. The first series of sections were stored in 10% formalin at room temperature and used for Nissl staining using thionin. The second series of sections were collected into tissue collection solution (30% ethylene glycerol, 25% glycerol in 0.05 M sodium phosphate buffer) and stored at −20 °C until staining with gold chloride for myelin^53^. Selected sections from WT and TG mice were used for immunohistochemistry. Free floating sections were stained for GFAP (1:1000, Millipore), Iba1 (1:500, Wako), WO2 (1:1000, Millipore) and DAPI (1:10000, Applichem). For permeabilization and blocking, buffer contained 1% BSA (Sigma), 0.2% fish gelatin (Sigma) and 0.1%Triton-X-100 (Sigma). Sections were incubated with primary antibodies diluted in the same solution overnight at 4°C. Host-matched secondary antibodies were applied the next day for 2 hours at room temperature. Sections were mounted with Aqua-Poly Mount (Polysciences). Imaging of the sections was performed using a Leica Thunder Slide Scanner at 20× magnification.

### 2.5. Code Availability

MATLAB source code for preprocessing and Monte-Carlo data inversion is freely available at https://github.com/maximeYon/MMD. The acquisition sequence is available upon reasonable request depending on Paravision versions.

## 3. Results

*ω*MD-MRI produces a family of microstructural parameters derived from generalized diffusion tensor distributions^54^. While the information is novel, it can be confusing for readers unfamiliar with the technique. Table 2 summarizes the interpretation of selected ωMD-MRI metrics and compares them with analogous metrics from commonly used techniques.

**Table 2.** General interpretations of select *ω*MD-MRI metrics.

| Metric | Interpretation |
| --- | --- |
| $E[D_{\text{iso}}]$ | Mean diffusivity, similar to ‘mean diffusivity’ (MD) in DTI |
| $E[D_{\Delta}^2]$ | Mean microscopic anisotropy of diffusivity. Unlike ‘fractional anisotropy’ (FA) in DTI, it is independent of fiber orientation dispersion. |
| $E[R_1]$ and $E[R_2]$ | Mean relaxation rates, associated with molecular interactions, chemical exchange and compartmental exchange<sup>57</sup> |
| $V[x]$ | Variance of any metric $x$, a reflection of the intravoxel heterogeneity, and/or uncertainty in the inversion which is related to the signal-to-noise ratio. |
| $\Delta_{\omega/2\pi}$ metrics | The linear rate of change of a metric with frequency, a measure of diffusion restriction. $\Delta_{\omega/2\pi}E[D_{\text{iso}}]$ is most sensitive to specific compartment sizes (“sweet spots”), which depend on compartment geometry, unrestricted diffusivity, and the frequency range. |
| Bin-resolved metrics | Metrics evaluated within only the specified bin. Bin 1 contains contributions from anisotropic water populations, typically associated with axons or neurites. Bin 2 contains contributions from more isotropic or permeable compartments. |

Specifically for the bin-resolved metrics, the categorization of water populations by their diffusive properties was first done in Pierpaoli et. al.^55^, where voxels in white matter, gray matter and cerebral spinal fluid (CSF) regions have distinguishable but coupled diffusivity-anisotropy characteristics, which was in part attributed to partial volume contamination. The binning in *ω* MD-MRI overcomes this by operating within the voxel, taking advantage of the solution distribution estimate. By separating the free-water fraction (bin 3)^56^ from bins 1 and 2, bin-resolved metrics provide tissue-specific measures: bin 1 represents the anisotropic tissue fraction, whereas bin 2 represents the isotropic tissue fraction.

### 3.1. ROI-averaged solution distribution projections

Figure 3 shows ROI-averaged distributions projected at the 10^th^ and 90^th^ percentile diffusion frequencies on to different axis combinations, including 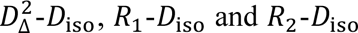, for the supra callosal cerebral white matter, cortical plate, and thalamus. The contribution of each subject to the plots was normalized.

For all three ROIs, there appears to be two populations, fast and slow diffusing, partially separable along *D*_iso_, most easily seen in the 1D distribution on top of each panel. The *R*_1_ and *R*_2_ variances in the fast diffusing population are larger than that of the lower *D*_iso_ value population, while the means are similar enough that the populations cannot be distinguished in the *R*_1_ and *R*_2_ 1-D projections. Components at the *R*_1_= 0 s^-1^ and *R*_2_= 100 s^-1^ edges^58^ are likely artifacts.

The distribution profile is more complex along 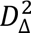, with two or more populations contained within bin 2 (shaded in yellow). There is also a sizeable population in bin 1 for the supra callosal cerebral white matter and thalamus, but too small to be observed in the cortical plate projections. At the maximal diffusion frequency, *ω*/2π = 247 Hz (right column in each subplot), the 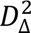 profile tends to simplify and concentrate towards 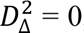, an expected behavior as diffusivities approaches the high-frequency isotropic diffusivity *D*_0_ (Eq 3) as frequency increases regardless of other properties.

The differences between the WT group and AD group are subtle. In all ROIs, the *D*_iso_ peak position of the fast diffusing population is higher in the AD group (in red). Also, the AD group has a higher *D*_iso_ variance, apparent from the lower peak heights. The *R*_2_ variance is higher in the AD group. Specifically, for the supra callosal cerebral white matter, the amplitude of the bin 1 peak is lower in the AD group

### 3.2 Visually apparent divergence in microstructure between AD and WT

Figures 4 shows group-averaged parameter maps for selected *ω*MD-MRI metrics from the WT mice and AD mice samples, as well as the percentage differences in the maps. Distinct features are visible in the percentage difference maps: in AD mice, E[*D*_iso_], E[*R*_1_], E[*R*_2_] and Δ*_ω_*_/2*π*_E[*D*_iso_] are generally higher, while 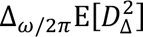 is more negative, particularly in the hippocampal formation. V[*D*_iso_] and V[*R*_2_] are also elevated in AD mice, mainly in the thalamus.

**Figure 4.**
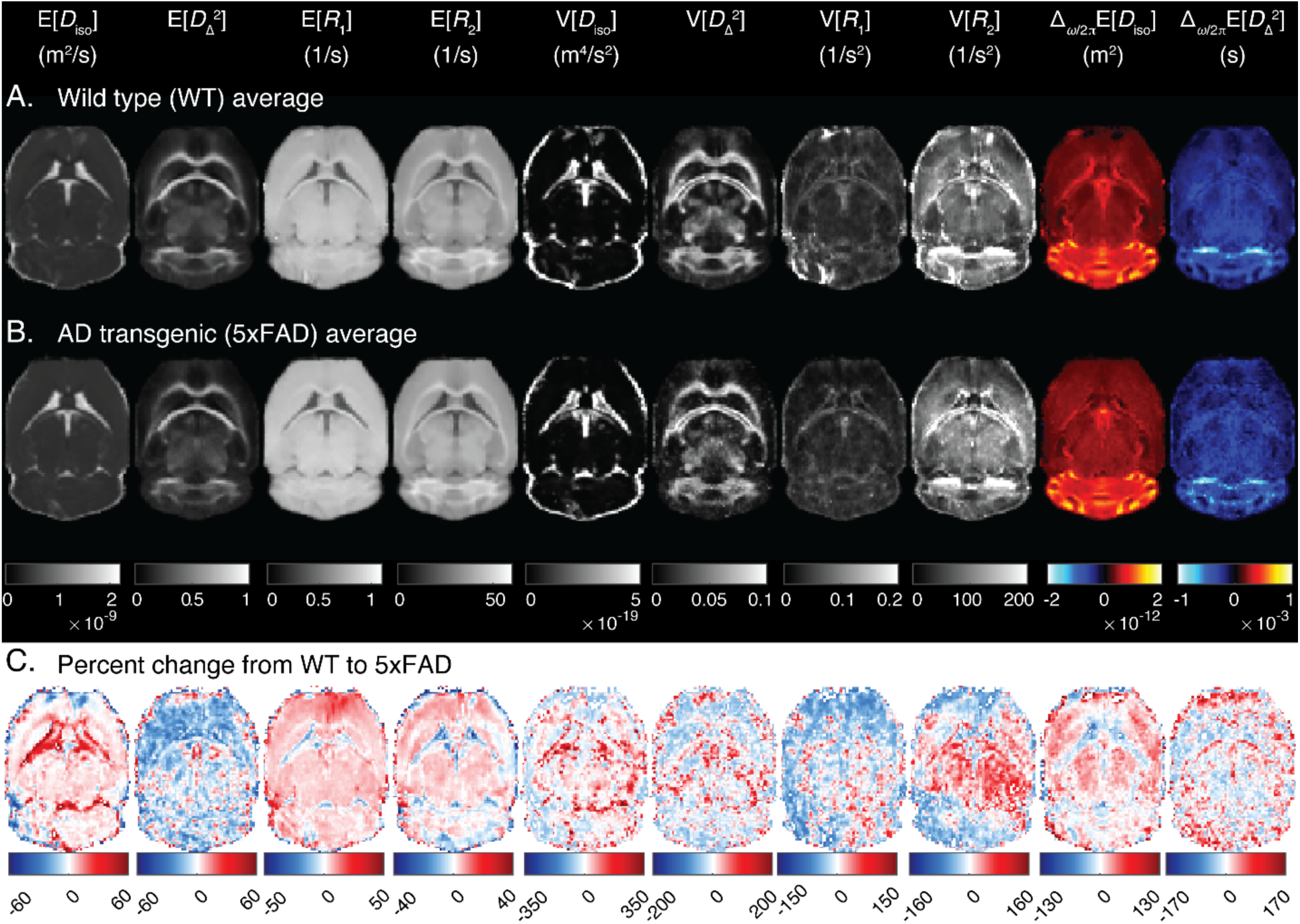
Group-averaged metric maps for selected metrics (including per-voxel means E[x] and variances V[x] evaluated at *ω*/2*π* = 36.6 Hz, and the rate of change with frequency Δ*ω*/2*π* of the per-voxel metrics) from the (A) WT and the (B) AD transgenic mice groups, and (C) the percent difference between WT and AD groups.

Fig. 5 shows the same parameters as in Fig. 4, only bin-resolved. For bin 1–resolved metrics, E[*D*_iso_]_bin1_, E[*R*_1_]_bin1_ and E[*R*_2_]_bin1_ showed a modest overall increase in the AD group, whereas 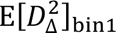 exhibited a slight decrease. Within the corpus callosum, 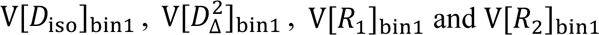 were lower in AD mice compared to controls. For bin 2–resolved metrics, similar trends were observed, with E[*D*_iso_]_bin2_, E[*R*_1_]_bin2_ and E[*R*_2_]_bin2_ displaying mild increases and 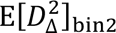 a slight decrease in AD mice. Additionally, 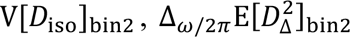 and Δ*_ω_*_/2*π*_E[*D*_iso_]_bin2_ showed more pronounced increases, while V[*R*_2_]_bin2_ was notably elevated in the thalamus of AD mice.

**Figure 5.**
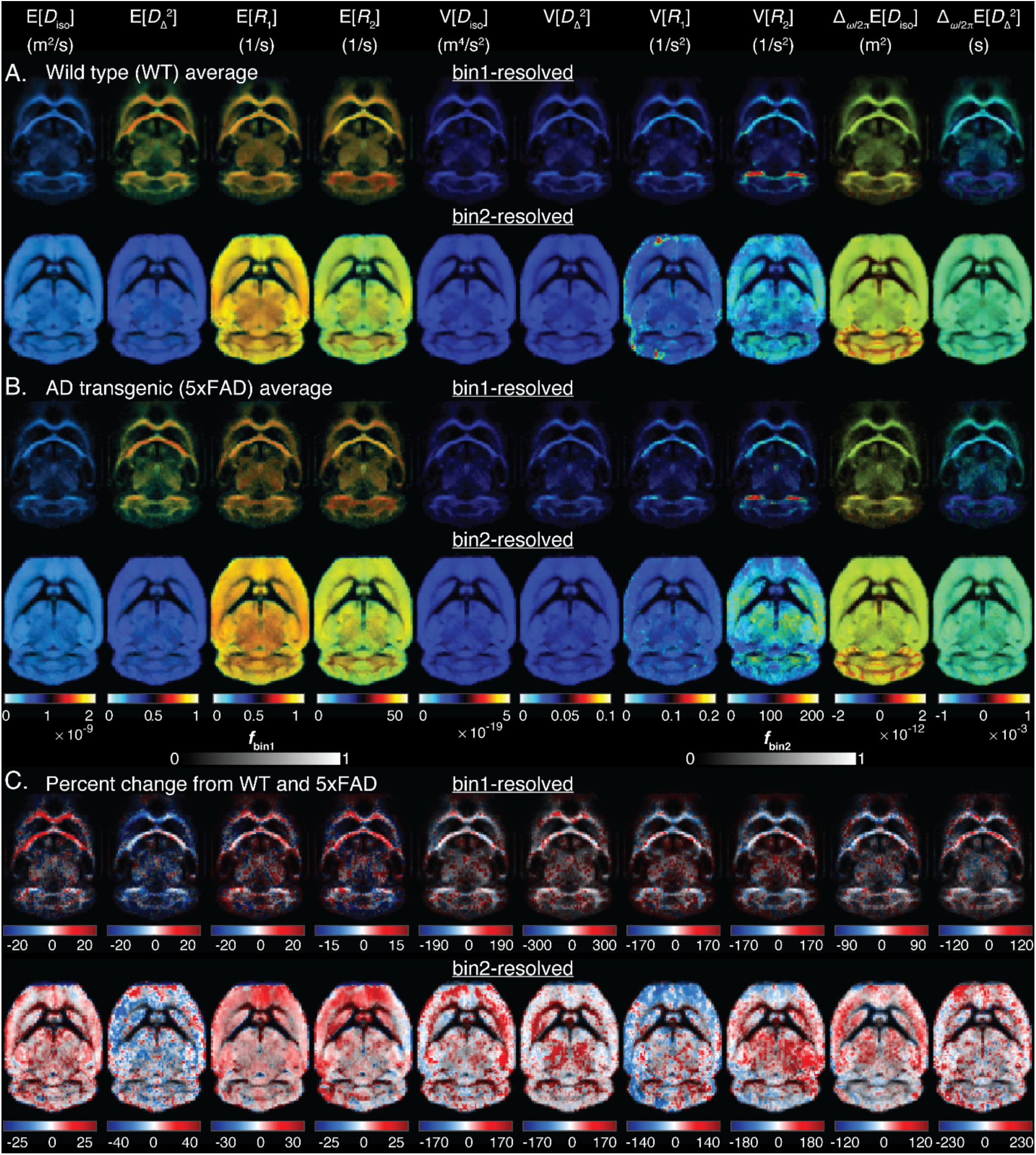
Group-averaged metric maps for selected bin-resolved metrics from the (A) WT and the (B) AD transgenic groups, and (C) the percent difference between WT and AD groups. On the top row of each subplot are metric maps resolved in bin 1 (anisotropic components, white matter dominant), and in the bottom row are the same but for bin 2 (isotropic components, gray matter dominant). These maps have been scaled according to the corresponding bin fractions.

For E[*D*_iso_]_bin2_, whereas the metric was generally higher in the AD group in most brain regions, in parts of the subicular complex, dentate gyrus and cortical plate the opposite was true (Fig. 6A). An increase in magnitude in percent difference was also observed for E[*R*_2_] and E[*R*_2_]_bin2_in the subicular complex (Fig. 6B).

**Figure 6.**
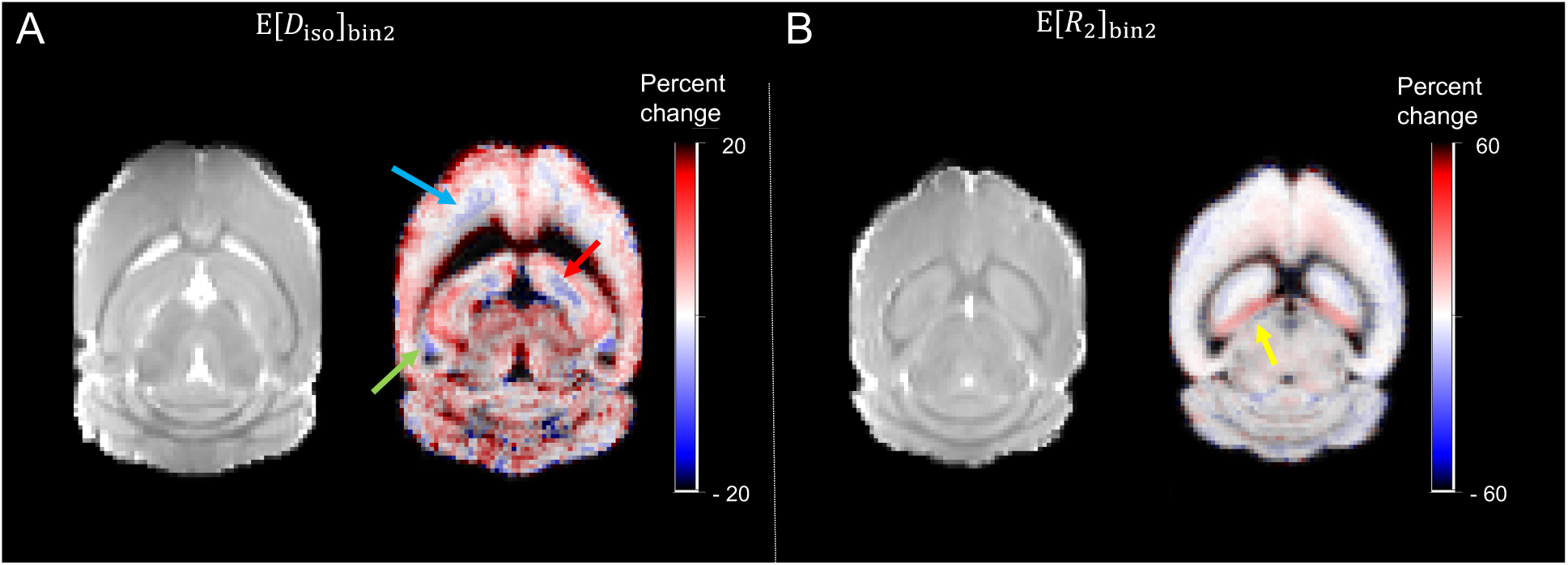
Group averaged metric percent change for (A) E[*D*_iso_]_bin2_ and (B) E[*R*_2_]_bin2_, accompanied by the *S*_0_ map. (A) and (B) show axial slices 0.6 mm and 1.2 mm dorsal relative to the location of the slice in Figs. 4 and 5 to display different anatomical regions. As elaborated in the results section of the main text, there are regions where the percent change of particular metric maps stand out from the overall trend, including regions in the subicular complex (green arrow), regions in the dentate gyrus (red arrow), regions in the isocortex (blue arrow), and subiculum (yellow arrow).

### 3.3. Increased sensitivity to diffusion frequency in AD

Figure 7 is a heatmap of effect sizes of metrics in each ROI distinguishing between WT and AD groups, obtained by linear regression of the Z-score adjusted results. Bin 3-resolved metrics, corresponding to properties of the free water fraction, have relatively lower repeatability^41^ as the bin 3 contribution in the defined ROIs are small (most ROIs have *f*_bin3_ < 0.05). Results for bin 3-resolved metrics are therefore presented in Supplementary Fig. 1.

**Figure 7.**
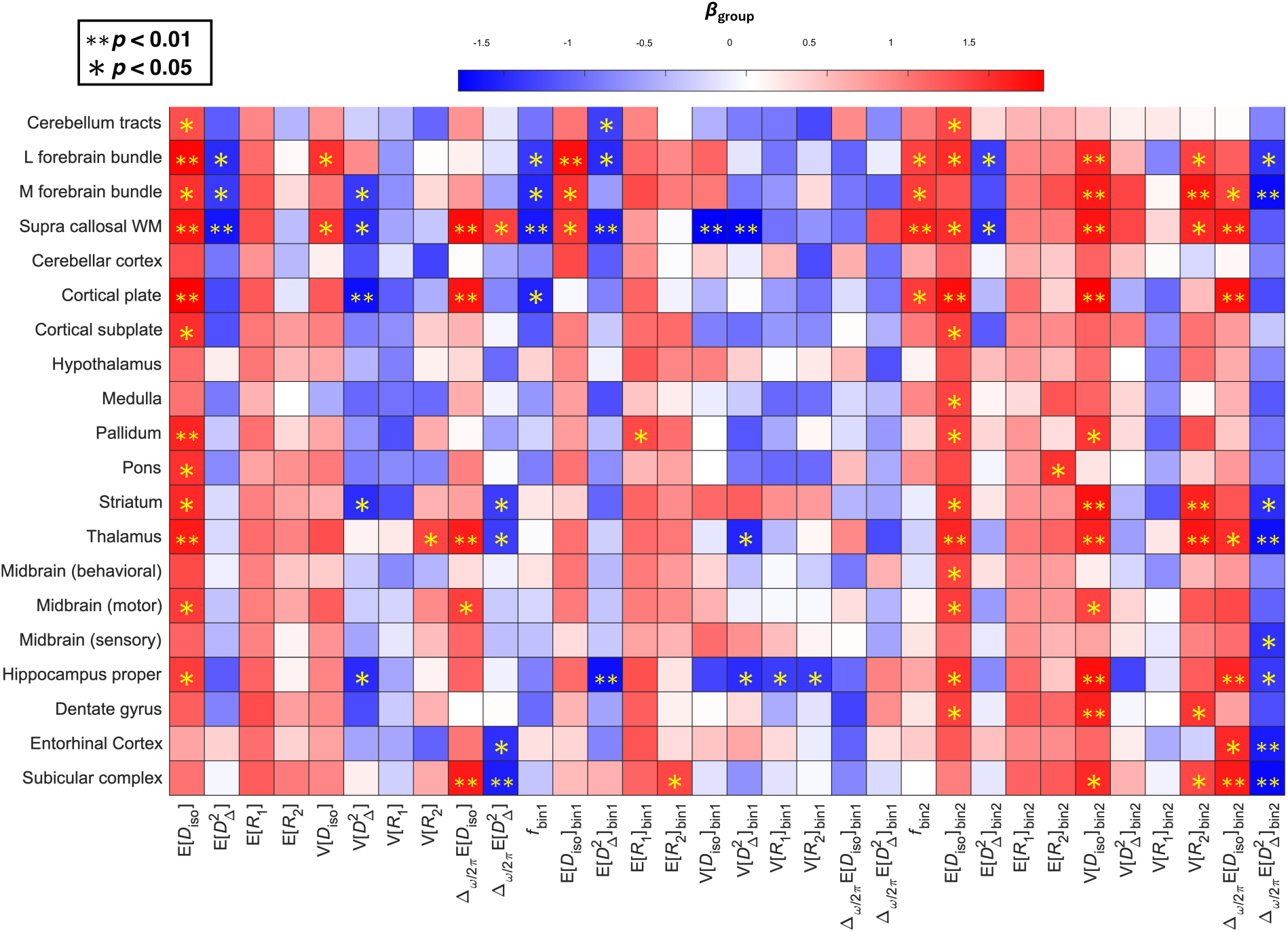
Heatmap of Z-score adjusted effect sizes (*β*_group_) of ROI-averaged *ω*MD-MRI parameters obtained by linear regression. Boxes in red are metrics that are higher in value in AD subjects, and blue indicates the opposite relationship. Statistical significances of *p* < 0.01 and *p* < 0.05 are notated as ∗∗ and ∗.

There were numerous statistically significant (*p* < 0.05) metrics that distinguish between the two groups in different ROIs. For most metrics, the difference between WT and AD mice was spatially consistent. For example, E[*D*_iso_] and E[*R*_1_] values were higher and 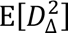 values were lower in AD mice in most ROIs. Diffusion frequency dependence metrics, 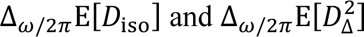 as well as their bin 2-resolved counterparts, also showed consistent trends across ROIs, with the former being higher and the latter being lower in AD mice. Note that 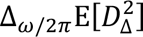 is a metric that generally has negative values, so the values are ‘more negative’ in AD mice.

A handful of metrics showed significant differences between the two groups across multiple ROIs, including E[*D*_iso_] (11 ROIs), E[*D*_iso_]_bin2_ (13 ROIs), V[*D*_iso_]_bin2_ (11 ROIs), V[*R*_2_]_bin2_ (7 ROIs), Δ*_ω_*_/2*π*_E[*D*_iso_]_bin2_ (6 ROIs) and 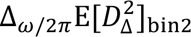 (8 ROIs). On the other hand, there were metrics that demonstrated spatial sensitivity to white matter fiber tracts, including 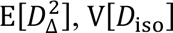, *f*_bin1_ and *f*_bin2_ (with the exception of the cortical plate), 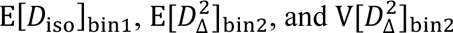.

The hippocampus is a region severely affected by AD, deserving more attention. Significant differences in the entorhinal cortex were observed only in the diffusion frequency parameters, specifically 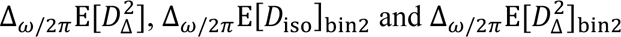. The diffusion frequency metrics also detected significant differences in the other hippocampus ROIs, namely hippocampal proper, dentate gyrus and subicular complex.

### 3.4. Histology

Immunohistochemical analysis of whole-brain sections revealed extensive amyloid pathology and associated gliosis in 5xFAD mice compared with WT controls (Fig. 8A). WO2-positive amyloid plaques were predominantly localized to the deep cortical layers, subiculum, hippocampus, and dentate gyrus, where they were accompanied by marked activation of both GFAP-positive astrocytes and Iba1-positive microglia. Although glial activation was strongest in the vicinity of plaques, increased astrocytic and microglial reactivity was also evident within plaque-free regions, including major white matter tracts such as the corpus callosum and cingulum, suggesting widespread neuroinflammatory alterations beyond the immediate sites of amyloid deposition (Fig. 8B). Histological assessment using Nissl and gold chloride staining showed preservation of overall cytoarchitecture and myelin organization with no evidence of massive neuronal loss or axonal degeneration in 5xFAD mice (Fig. 8C). Nevertheless, local structural distortion was observed around plaques, including clustering of glial cells and axons bending around amyloid deposits.

**Figure 8.**
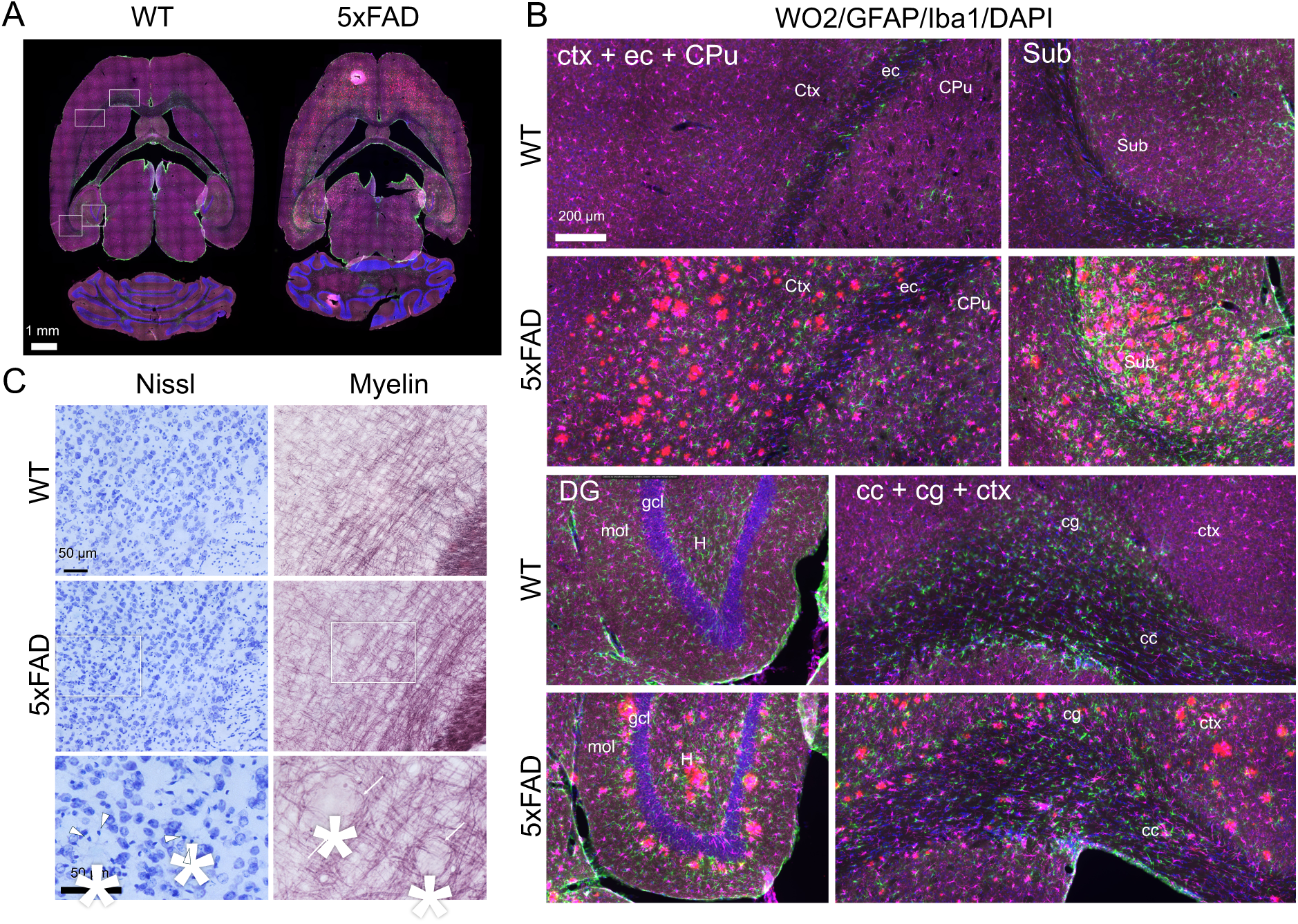
Histological comparison between a wild-type (WT) and a transgenic (5xFAD) mouse brain. (A) Whole-section images of WT and 5xFAD mouse brain sections showing the four imaging channels: GFAP (green), Iba1 (magenta), WO2 (red), and DAPI (blue). Scale bar: 1 mm. (B) Closeup views of selected brain regions from the image shown in A, including the cortex (ctx), external capsule (ec), caudate putamen (CPu), subiculum (Sub), dentate gyrus (DG), corpus callosum (cc), and cingulum (cg). Plaque accumulation was most prominent in the deep cortical layers, subiculum, hippocampus, and dentate gyrus. Activation of both astrocytes and microglia was strongest at plaque sites but was also observed in regions devoid of plaques, including white matter tracts. Scale bar: 200 µm. (C) Representative photomicrographs of two consecutive sections to the section shown in A, stained with Nissl staining for cytoarchitecture and gold chloride for myelin, respectively. Images depict the deep cortical layers of the primary somatosensory cortex. White asterisks indicate plaque locations, white arrowheads indicate glial cells around the plaque and white arrows point at axons bending around plaques. Comparing WT and 5xFAD, no massive neuronal or axonal degeneration was observed. Scale bar: 50 µm.

## 4. Discussion

The present study demonstrates that ωMD-MRI detects widespread, regionally heterogeneous microstructural differences between 5xFAD and WT mouse brains that are not fully captured by conventional diffusion-derived measures. The most spatially consistent group differences were observed in measurements sensitive to diffusion-frequency dependence, intravoxel heterogeneity, and bin-resolved water components, with particularly prominent abnormalities in the hippocampal formation and major cerebral white matter tracts. Histological analyses revealed extensive amyloid deposition accompanied by marked astrocytic and microglial activation, including inflammatory changes in white matter regions with limited plaque burden. Despite these pathological changes, overall cytoarchitecture and myelin organization remained preserved, with no evidence of widespread neuronal loss or axonal degeneration, although localized tissue distortion was evident around plaques. Together, these findings indicate that the ωMD-MRI abnormalities observed in the examined 5xFAD mice reflect a pathological state characterized by amyloid deposition, gliosis, and localized tissue remodeling rather than widespread tissue destruction.

A key finding of this study is that the pathological state observed in the examined 5xFAD mice is associated with substantially greater microstructural heterogeneity than in WT animals. Histological examination demonstrated that amyloid plaques, reactive astrocytes, activated microglia, localized tissue distortion, and relatively preserved parenchyma coexist within the same anatomical regions, creating a complex mosaic of microscopic environments. Conventional diffusion MRI necessarily averages these diverse water environments into a single apparent diffusion coefficient per voxel, limiting the ability to distinguish their individual contributions to the measured signal. In contrast, ωMD-MRI characterizes multidimensional distributions of water components with distinct diffusion and relaxation properties, providing a more comprehensive description of tissue microstructure at the voxel level. The widespread differences observed in heterogeneity-sensitive measurements, together with alterations in the bin-resolved analysis, are consistent with this interpretation and suggest that the potential benefit of ωMD-MRI may extend beyond increased sensitivity to providing additional information about underlying microstructural complexity that may be obscured in conventional diffusion MRI measurements.

The histological findings provide important biological context for interpreting the multidimensional imaging measurements. Amyloid plaques were abundant throughout the cortex, hippocampus, dentate gyrus, and subiculum and were accompanied by pronounced astrocytic and microglial activation. Inflammatory changes were also evident in white matter regions with relatively low plaque burden, indicating that tissue remodeling extends beyond the immediate vicinity of amyloid deposition. Despite these pathological alterations, Nissl and gold chloride staining demonstrated preservation of overall cytoarchitecture and myelin organization, with only localized distortion surrounding individual plaques. This preservation should not be interpreted as evidence that myelin or axonal pathology is absent, because subtler alterations in tissue integrity may not be detectable with the histological methods used. These observations suggest that the imaging abnormalities reflect the combined influence of multiple coexisting pathological features, including amyloid deposition, reactive gliosis, neurite remodeling, and alterations of the extracellular microenvironment, rather than any single biological process. Consequently, individual multidimensional diffusion measurements should not be interpreted as specific markers of individual pathological substrates. Instead, they provide complementary descriptions of the underlying microstructural architecture^59^ of diseased tissue arising from the simultaneous presence of multiple interacting pathological processes.

Measurements incorporating diffusion-frequency dependence were among the most consistently altered throughout the 5xFAD brain, particularly within the hippocampal formation and associated cortical regions. By probing progressively shorter effective diffusion distances, these measurements are sensitive to changes in tissue organization occurring over microscopic length scales, where water motion is influenced by cellular membranes, neurites, glial processes, and other structural barriers. The broad spatial distribution of these abnormalities suggests that the pathological changes present in the examined 5xFAD mice involve alterations in tissue architecture that extend beyond changes detectable by conventional diffusion-derived parameters alone. Importantly, these observations should not be interpreted as evidence that diffusion-frequency-dependent measurements reflect a specific pathological feature, such as amyloid plaques or reactive gliosis in isolation. Rather, they indicate sensitivity to the cumulative effects of microscopic tissue remodeling arising from the coexistence of multiple pathological processes. In this context, the value of ωMD-MRI is not simply the generation of additional quantitative measurements, but its ability to interrogate tissue organization across multiple diffusion length scales while simultaneously resolving multiple water components within the same imaging voxel.

A further important observation was that the multidimensional measurements revealed the spatial complexity of pathological remodeling within individual anatomical structures rather than simply identifying affected regions. This was particularly evident in the cerebral cortex and hippocampal formation. Within the cortex, percentage-change maps of the heterogeneity- and diffusion-frequency-sensitive measurements qualitatively mirrored the laminar distribution of WO2-positive amyloid pathology (Fig. 9), with the largest changes of imaging metrics occurring in the deep cortical layers where plaque burden was greatest. Likewise, although the hippocampal formation consistently exhibited some of the largest group differences, individual subregions displayed distinct imaging signatures, including localized reversals in diffusivity-related measurements within portions of the dentate gyrus and subicular complex (Fig. 6). Together, these observations indicate that the pathological state in the examined 5xFAD mice is spatially heterogeneous across multiple anatomical scales, extending from cortical layers to hippocampal subregions. Although the qualitative correspondence between imaging and histology does not establish biological specificity, it demonstrates that tissue remodeling is not uniformly distributed within affected brain structures. Future quantitative voxel-wise MRI–histology correlation studies will be important for defining the biological basis of these localized imaging signatures.

**Figure 9.**
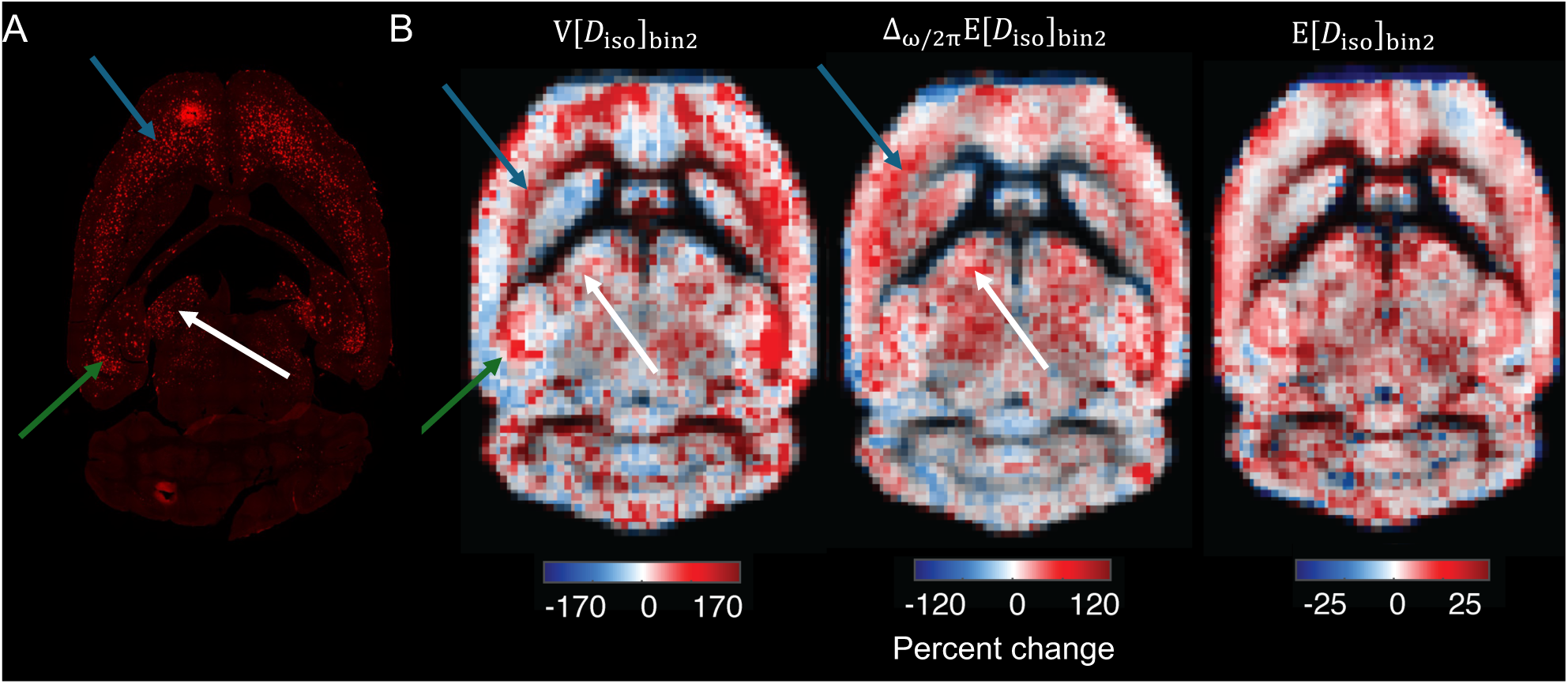
(A) WO2 staining of amyloid beta plaques and (B) percent change from WT group to AD group of MRI metric maps of V[*D*_iso_]_bin2_, Δ*_ω_*_/2*π*_E[*D*_iso_]_bin2_ and E[*D*^2^]_bin2_. Blue, green and white arrows indicate inner layers of the cortical plate, subiculum, and thalamus respectively. V[*D*_iso_]_bin2_ resembles the WO2 staining in all 3 regions, while Δ*_ω_*_/2*π*_E[*D*_iso_]_bin2_ resembles only the cortical plate and thalamus.

The white matter findings further emphasize the widespread nature of the pathological changes detected by the multidimensional diffusion measurements. Although AD has traditionally been viewed as a predominantly gray matter disease, increasing evidence indicates that white matter alterations are an important component of AD pathology in both patients^60,61^ and transgenic mouse models^62^. In the present study, significant abnormalities were observed throughout the major cerebral white matter tracts, whereas cerebellar pathways remained comparatively unaffected, paralleling the known regional distribution of pathology in the 5xFAD model. Histological analyses also demonstrated inflammatory activation within white matter despite relatively limited plaque deposition, suggesting that tissue remodeling is not restricted to plaque-rich regions. Interpretation of the isotropic bin-resolved measurements within white matter should nevertheless be approached with caution, as the dimensions of many fiber tracts approach the imaging voxel size, making partial-volume effects difficult to avoid. Consequently, the anisotropic bin-resolved measurements likely provide the more robust characterization of white matter architecture. Although the observed imaging abnormalities are consistent with subtle alterations in tissue architecture, the present data do not distinguish among potential contributions from inflammatory remodeling, changes in axonal organization, myelin ultrastructure, or other microscopic features of white matter. The absence of overt myelin disruption should therefore not be interpreted as evidence against myelin involvement in AD, particularly given evidence of mechanistic interactions between myelin integrity, amyloid pathology, and associated inflammatory responses^3^.

The present findings are broadly consistent with previous diffusion MRI studies of the 5xFAD model while extending the characterization of disease-associated microstructural alterations. Previous studies consistently reported abnormalities within cortical and cerebral white matter regions, most commonly manifested as changes in diffusivity and reductions in fractional anisotropy, supporting the presence of widespread tissue abnormalities in both mice and patients with AD^15–17^. These studies demonstrated that diffusion MRI is sensitive to disease-associated tissue changes but also highlighted the limitations of measurements that average multiple pathological processes within the MRI voxels. The present study reproduced many of these established regional abnormalities while demonstrating that multidimensional diffusion encoding provides additional insight into their underlying microstructural complexity. In particular, measurements sensitive to diffusion-frequency dependence, microstructural heterogeneity, and bin-resolved water components consistently identified abnormalities throughout the hippocampal formation, where previous diffusion MRI studies have often reported more variable findings. Taken together, these observations support the view that the microstructural alterations associated with AD are inherently multidimensional, reflecting the coexistence of multiple tissue environments that cannot be fully represented by a single voxel-averaged diffusion value.

Several limitations should be considered when interpreting these findings. First, this was a cross-sectional ex vivo study performed in a single transgenic mouse model at a single disease stage, limiting conclusions regarding the temporal evolution of the observed microstructural alterations. Second, the relatively small sample size and inclusion of only female mice may limit the generalizability of the findings. Third, although the qualitative correspondence between multidimensional imaging measurements and histological abnormalities supports their biological relevance, the present study was not designed to establish direct voxel-wise relationships between individual imaging metrics and specific pathological substrates. Such analyses will require dedicated image-registration strategies and quantitative histological assessment. Fourth, the 5xFAD model primarily reflects amyloid pathology and does not recapitulate the full spectrum of human AD. In particular, the substantially lower white matter and myelin content of the mouse brain may limit the extent to which this model captures the contribution of myelin pathology and long-range white matter degeneration to human disease, and this should be considered when interpreting the absence of overt myelin or axonal degeneration despite extensive amyloid pathology. Finally, although the ex vivo experimental design enabled high spatial resolution and comprehensive multidimensional diffusion encoding, future in vivo studies, employing feasible acquisition protocols^63–65^, will be needed to determine the feasibility of translating these measurements to longitudinal studies and, ultimately, clinical applications.

In conclusion, the present findings demonstrate that AD pathology in the 5xFAD mouse brain is associated with complex and spatially heterogeneous tissue remodeling arising from multiple coexisting microstructural environments. This complexity cannot be adequately represented by conventional diffusion MRI measurements that average these contributions into a single signal. By resolving complementary features of tissue heterogeneity across water environments and diffusion length scales, and relating them to histopathological abnormalities, ωMD-MRI provides a more comprehensive framework for characterizing the biological complexity of AD. Although the biological basis of individual measurements and their in vivo translatability require further validation, this approach may ultimately support more informative studies of disease progression and therapeutic response.

## Acknowledgements

This research was supported by the Intramural Research Program of the National Institutes of Health (NIH). The contributions of the NIH author(s) are considered Works of the United States Government. The findings and conclusions presented in this paper are those of the author(s) and do not necessarily reflect the views of the NIH or the U.S. Department of Health and Human Services. This work was supported by the Swedish Research Council (Vetenskapsrådet; grant no. 220222-04422_VR), the Research Council of Finland (Academy Project Funding #361370, and Flagship of Advanced Mathematics for Sensing Imaging and Modelling (FAME) #358944).

The authors thank Murat Bilget and Sridhar Kandala at NIA/NIH for their advice on the statistical analysis of the results, and to Maarit Pulkkinen at UEF for her support with animal handling and histology.

## Disclosure Statement

The authors declare no conflicts of interest.

## Notes

### Competing Interest Statement

The authors have declared no competing interest.

