## Supplementary for "Multidimensional diffusion MRI reveals heterogeneous microstructural remodeling associated with amyloid pathology"

| Sample | Weight (g) | Sex | Age at perfusion (days) | Time between perfusion and scan (days) |
|---|---|---|---|---|
| WT1 | 24 | F | 258 | 270 |
| WT2 | 24 | F | 258 | 266 |
| WT3 | 24 | F | 258 | 333 |
| WT4 | 25 | F | 258 | 275 |
| AD1 | 21 | F | 234 | 340 |
| AD2 | 24 | F | 234 | 228 |
| AD3 | 22 | F | 234 | 341 |
| AD4 | 21 | F | 234 | 235 |

**Supp Table 1.** details of subjects, dates of perfusion and dates of chosen scans

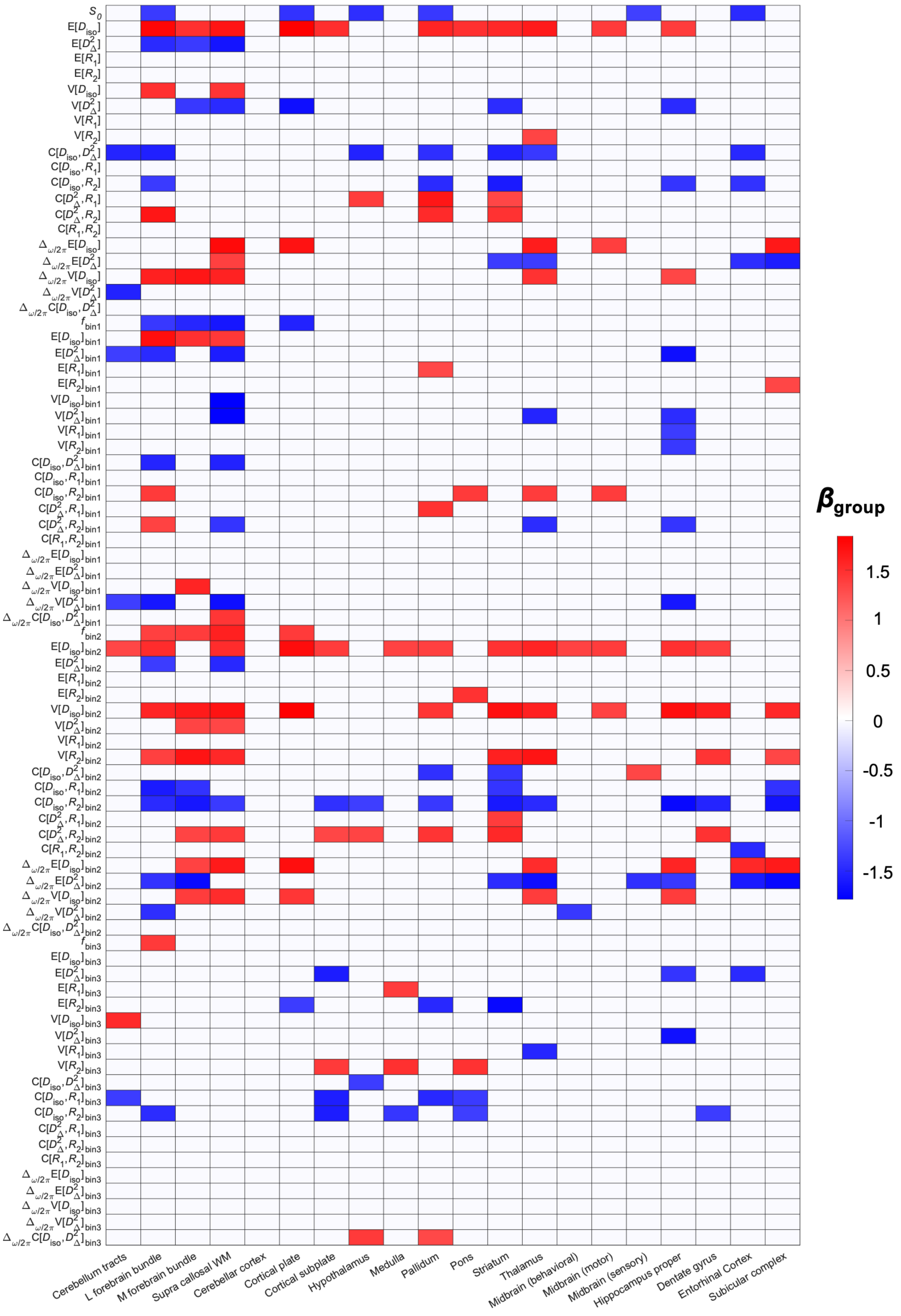


**Supp Fig 1.** Heatmap of linear regression effect sizes $\beta_{\text{group}}$ with all metrics. Only entries where the difference is significant ($p$<0.05) are colored for viewability. Along the x-axis are the metrics, in the order of $S_0$, $\mathrm{E}[x]$, $\mathrm{V}[x]$, $\mathrm{C}[x, y]$, $\Delta_{\omega/2\pi}$ metrics, and then corresponding bin 1-, 2- and 3-resolved metrics. Along the y-axis are the ROIs.
